# Integrative omics data analysis uncovers biomarker genes and potential candidate drugs for G3 medulloblastoma

**DOI:** 10.1101/2020.06.19.122515

**Authors:** Loreina Guo, Kendall Cornick, Vincent Xu, Tianyao Hao, Ma. Xenia G. Ilagan, William Buchser, Joshua B. Rubin, Fuhai Li

## Abstract

Medulloblastoma (MB) is the most common malignant brain tumor in infants and children. Four molecular subtypes of MB are recognized: WNT, SHH, Group 3 (G3), and Group 4 (G4). Compared with WNT and SHH subtypes, G3 MBs exhibit significantly worse outcomes and higher metastatic rates, and there is no effective treatment yet. Moreover, G3 and G4 MBs are much more common in boys than girls, i.e., sex bias, which also plays important roles in cancer prognosis and drug response. However, the molecular mechanism of G3 remains unclear, and there are no well-identified biomarker genes associated with these phenotypes, i.e., worse survival rate, higher metastasis rate, and sex bias. In this exploratory study, we aim to identify potential biomarkers associated with the three phenotypes using integrative analysis of gene expression, methylation and copy number variation datasets. In the results, we identified a set of biomarker genes and linked them into a network signature. The network signature showed better performance in the separation of G3 MB patients into subtypes with a significant difference in terms of the three phenotypes. To identify potentially effective drugs for G3 MBs, a set of drugs with diverse targets were prioritized, which can potentially inhibit the network signature. These drugs or combinations thereof might be effective for G3 treatment.

## 1. Introduction

Medulloblastoma (MB) is the most common malignant brain tumor in infants and children occurring predominantly between the ages of 3 and 8 years old^1^. Four molecular subtypes of MB are currently recognized: WNT, SHH, Group 3 (G3), and Group 4 (G4) ^2,3,4^. G3 and G4 occur most frequently accounting for > 60% of MB cases while exhibiting significantly worse outcomes than WNT and SHH^5,6^. G3 MBs predominantly affect infants and children^1,5^ and have the worst 5-year survival rate of only 50% with a 50% metastasis rate at the time of diagnosis^1,3,4^. Moreover, G3 and G4 MBs have a much higher prevalence rate in boys than girls. In recently studies, sex of glioblastoma multiforme (GBM) patients was found to be correlated with prognosis and responses to different treatments^7,8^. These results indicated that sex of cancer patients might be an important factor for designing personalized treatment. Though the molecular subtyping analysis of MBs is a kind of biomarker analysis for poor survival, the molecular mechanisms and biomarker genes underlying G3 MBs are less clear compared with WNT and SHH MBs^2,3^. To date, no specific integrative multi-omics data analysis has been done to uncover the biomarkers associated with all the three phenotypes simultaneously, i.e., the worse survival rate, higher metastasis rate, and sex prevalence difference phenotypes in G3 MBs, which play important roles in understanding the molecular mechanism and developing personalized precision treatment regimen. Therefore, there is a significant need to identify the biomarker genes associated with the three phenotypes in G3 MBs specifically.

There is no effective targeted therapy that has been approved by the FDA for G3 MBs^5^, though some drugs and compounds have been reported in the literature. Also, G3 MBs is the worst prognostic group. The current treatment, including surgery, radiation and chemotherapy, is aggressive and toxic. Thus, novel effective treatments for children suffering with G3 medulloblastoma are urgently needed. It is important to uncover the signaling mechanisms and biomarkers that are associated with the low survival rate, high metastasis rate, and higher prevalence in boys, and consequently discover novel and effective drugs and drug combinations perturbing these signaling biomarkers as candidates for G3 MB treatment. On the other hand, there are diverse data resources that can be used to discover and repurpose potential targeted therapies for cancer treatment^9^. For example, the connectivity map^10,11^ (CMAP) generated a large number of reverse gene signatures in many cancer cells before and after drug treatment with 2837 different drugs and compounds. The Broad Institute Cancer Cell Line Encyclopedia (CCLE)^12^ and Genomics of Drug Sensitivity in Cancer (GDSC)^13,14^ datasets associated the genomics biomarkers of cancer cells with the responses to hundreds of drugs. Moreover, the drug targets^18,19^ can also be used to discover drugs for cancer. Also, the genomics data, e.g., disease associated genes^22^, differentially expressed gene sets^23,24^, GO terms^25^, and network analysis^26,27^, are often combined with aforementioned information to reposition drugs for cancer.

In this study, we aim to identify the informative biomarkers associated with the three phenotypes in G3 MBs by integrating the gene expression data, methylation data and copy number/SNP data. The resulting biomarkers were then connected to form a signaling network by using the BioGRID^30^ protein-protein interaction data. Consequently, the signaling networks were used as network signatures to discover drug candidates that can potentially inhibit these biomarkers by using drug-target information derived from DrugBank and reverse gene signature data available from CMAP data. To investigate the potential mechanisms of hundreds of the drugs selected, we divided drugs into clusters by using the drug target information and affinity propagation (AP) clustering. Importantly, our approach rediscovered many drugs that have been reported in the literature to be related to MB treatment, thereby validating our analysis approach.

## 2. Method

### 2.1 Gene expression, Methylation, SNP data and clinical data of MB

In the present study, expression, methylation, copy number, and clinical profiles of MB samples were downloaded from the Gene Expression Omnibus (GEO) (see **Table I**). Expression data were downloaded from **GSE85217**. Methylation data were downloaded from GSE85212^31^. There are 763 total samples; 144 are Group3, 326 in Group4, 70 in SHH, and 223 in WNT. Among these samples, Group4 has 92 females and 216 males, Group3 has 38 females and 99 males, WNT has 35 females and 29 males, and SHH has 82 females and 128 males. SNP data were downloaded from GSE37384^32^. The other samples do not have the gender information available. There are 1097 samples and 484 of them are matched with expression and methylation data. There are 21641 probes in the expression and 321172 probes in the methylation data.

**Table I:**
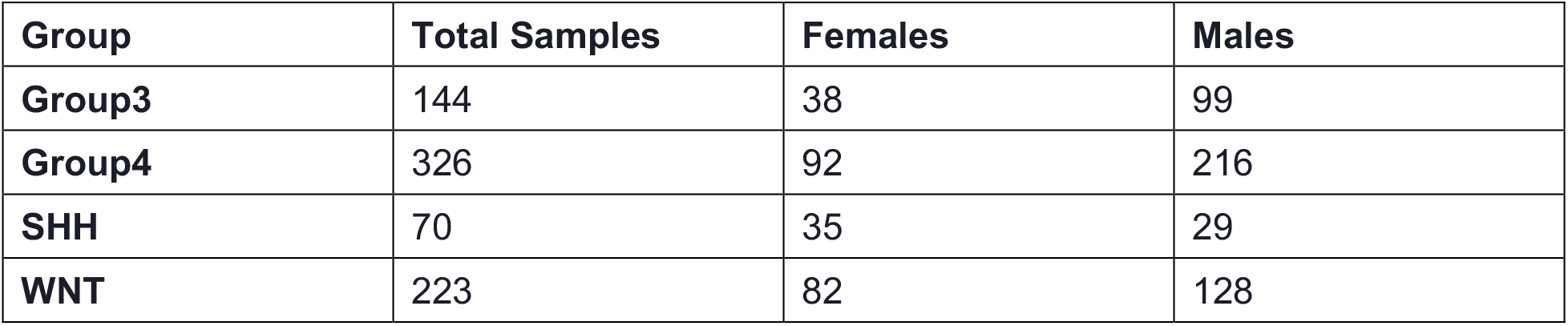
Number of samples in each MB subtype.

| Group | Total Samples | Females | Males |
| --- | --- | --- | --- |
| Group3 | 144 | 38 | 99 |
| Group4 | 326 | 92 | 216 |
| SHH | 70 | 35 | 29 |
| WNT | 223 | 82 | 128 |

### 2.2 Data Analysis and Integration

#### 2.2.1 Expression data analysis

The fold change and p-value of individual genes (probes) were obtained by comparing the mean gene expression data and conducting a student t-test between Group3 versus WNT subtypes. Up-regulated genes were then picked with a fold change >= 1.5 and p-value <= 0.05. Transcription Factors (TFs) and associated target genes were downloaded from MSigDB database^33,34^. We identified the activated TFs using the gene set enrichment analysis by using the Fisher exact test and comparing the number of up-regulated target genes of each individual TFs. Activated TFs were then selected with a p-value <= 0.00001.

#### 2.2.2 Methylation data analysis

The fold change and p-value of individual genes (probes) were obtained by comparing the mean methylation data and conducting a student t-test between Group3 versus WNT subtypes. The top 5000 hypo-methylated probes were selected (with a p-value less than 0.05). To reduce the number of biomarker genes from methylation data, only the hypo-methylated genes were kept, which also appeared in the list of selected up-regulated gene**s**, or the activated TFs, or has interactions with the up-regulated gene**s** or activated TFs in the BioGRID^30^ protein-protein interaction network.

#### 2.2.3 Copy number data analysis

Raw SNP (Affymetrix 6) array data were analyzed using ‘rawcopy’ R library^35^. The copy number of individual genes were obtained from the segmentation file generated by ‘rawcopy’. The normal control samples (Affymetrix 6) were provided by ‘rawcopy’. The amplification peaks were identified with a threshold of an average log2 ratio larger than 1.5.

#### 2.2.4 Phenotype specific biomarker selection using gene expression and methylation data

To identify genes associated with longer survival, Pearson correlation analysis between survival time (for shorter and longer survival) and the gene expression fold change and the methylation fold change of individual genes was conducted for G3 respectively. For the metastasis phenotype, MB samples in G3 were divided into non-metastasis and metastasis subgroups, and then the differentially expressed and methylated genes between the metastasis and non-metastasis groups were identified based on the p-values (<= 0.05) from the student t test. The same analysis was conducted to identify sex-specific genes. The selected biomarker genes were then filtered by intersecting the up-regulated genes and the hypo-methylated genes in G3 compared with the WNT MB subtype.

#### 2.2.5 Phenotype-specific biomarker selection using SNP/copy number data

MB samples in G3 were divided into male and female, metastasis and non-metastasis, and survival time longer than average and the survival time shorter than average groups respectively. The frequency of genes with amplified copy number, appeared in the MB patients, was calculated for each of the above groups. Genes that had a higher frequency in the male, metastasis, and longer survival groups compared to the female, non-metastasis and shorter survival groups respectively, were identified as biomarkers for each specific phenotype.

#### 2.2.6 Biomarker Network Construction

The selected phenotype-specific biomarkers, obtained from gene expression, methylation, SNP, were then connected into a signaling network using the BioGRID protein-protein interaction network as the background graph. Specifically, the ‘shortest paths’ (which is defined as the shortest protein-protein interactions/paths linking the two given biomarker proteins in the BioGrid network) among the selected biomarkers on the BioGRID network were obtained using the ‘igraph’ ^36^ R library to link all the biomarkers into a connected signaling network. The biomarker genes with the shortest-path greater than 5 were not included into the signaling network.

#### 2.2.7 Prioritization of drug targeting on the signaling network

The drugs and investigational agents were selected as followed. First, FDA-approved drugs, whose targets are on the signaling network, were selected. Second, the signaling network genes were used as gene signatures in the Connectivity Map (CMAP)^11^ database (clue.io) to predict drugs that can potentially inhibit these genes. Drugs with ‘summary score’ from CMAP less than ‘-80’ were selected as candidates for G3/G4 treatment.

#### 2.2.8 Drug clustering analysis

To understand the potential mechanisms of these drugs, we clustered drugs into sub-groups based on their targets. First, drug-drug similarity was evaluated by the Jaccard index of drug targets: 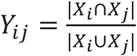. Then, the drug-drug distance matrix was obtained as a similarity matrix. Then, the propagation(AP) clustering algorithm^37,38^ was employed to cluster drugs into sub-clusters.

## 3. Results

### 3.1 Biomarkers genes and GO analysis for G3 MBs

A number of genes were selected for associated with the three phenotypes, e.g., male vs. female, metastasis vs. non-metastasis, and longer survival rate vs. shorter survival rate, in G3 MBs. Specifically, for G3 MBs, 66 biomarkers (gene expression: 28 + methylation: 21 + copy number: 17) were identified for male vs. female difference; 75 biomarkers (gene expression: 28 + methylation: 26 + copy number: 21) were identified for metastasis vs. non-metastasis difference; and 56 biomarkers (gene expression: 24 + methylation: 26 + copy number: 6) were identified for longer survival rate vs shorter survival rate difference.

Using the BioGRID protein-protein interaction network, we connected all of the selected biomarker genes into a signaling network by linking these biomarkers using the shortest paths in the BioGRID network. **Fig. 1** shows the integrative signaling networks of G3 MBs. In summary, there are 128 genes in the G3 integrative signaling network. Specifically in G3 MBs, the biomarkers MYC, COL3A1, COL1A1, COL1A2, PPARG, TGFBI have been reported to be important in G3 MBs ^39^. MYC amplification or overexpression is believed as one of major biomarkers of G3 MBs. Moreover, MB patients with MYC gene amplification tend to have a worse survival rate than other patients^40^. COL3A1, COL1A1, and COL1A2 are also important biomarkers and play a pre-growth role ^41^. Elevated levels of PPARG have been found in the clinical samples of MB^42^. Valproic acid, which is a drug that inhibits PPARG, has been reported to inhibit cell proliferations and induce MB cellular senescence and differentiation^43^. In addition, TGFB1 is another important and over-expressed gene, and the Hyaluronidase that targets TGFB1 has been reported as a potential MB treatment, but requires further experimental validation in vivo^44–47^. Also, Nicardipine has been reported as an inhibitor of the voltage-gated calcium channel CACNA1C^48,49^, which has been reported to be differentially expressed in MB, and can be one of the novel targets to treat cancer^50^. EGFR also has been found to be a potential target alone or combined with other targets to treat MB^51^. For example, there is potential for Cetuximab (EGFR inhibitor) to be combined with BIM-23127 (the peptidic neuromedin B receptor antagonist) to inhibit EGFR to induce the death of MB cells^52^.

**Figure 1.**
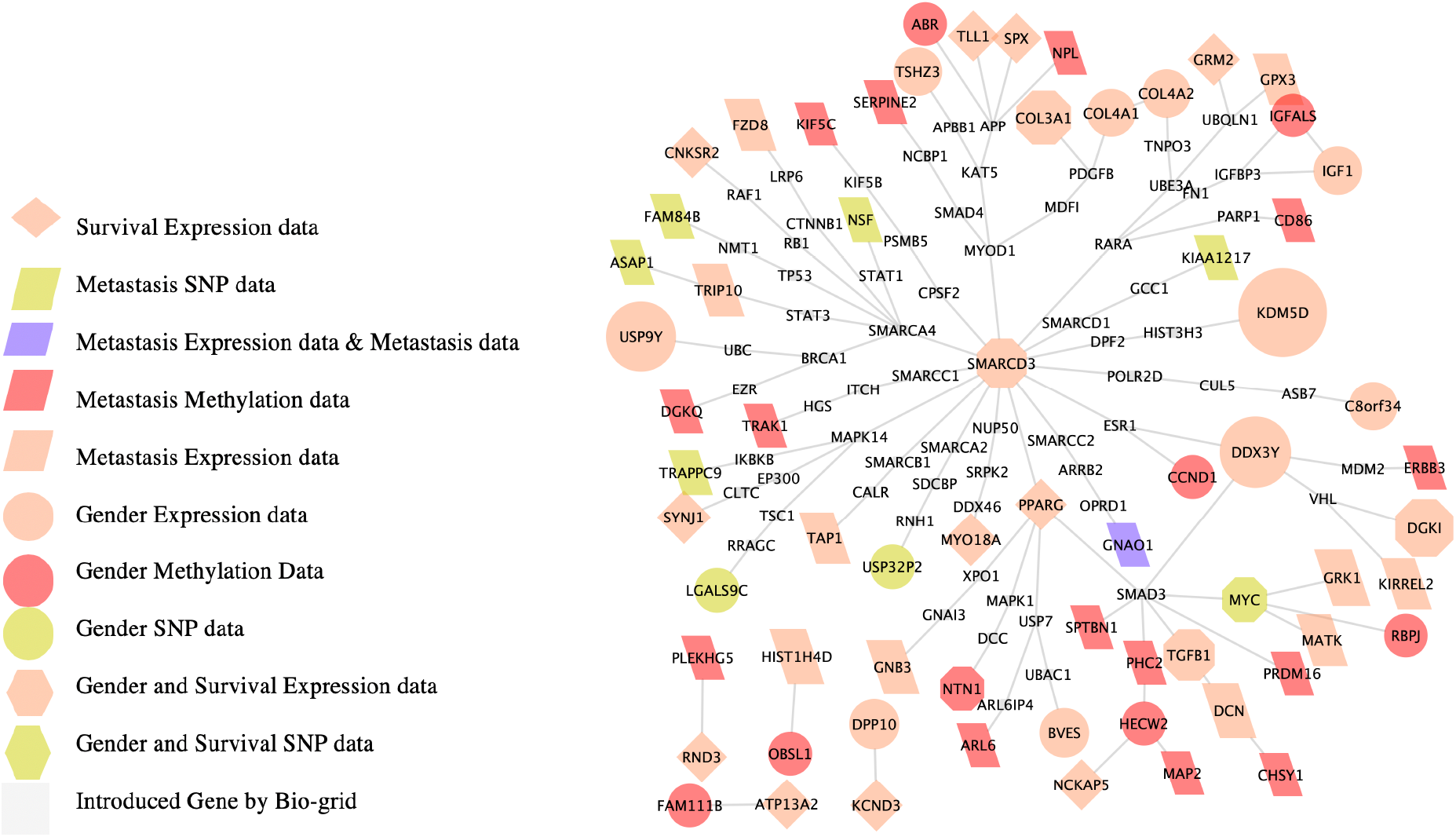
The signaling network consists of 52 biomarker genes in G3 MBs. Different shape and color indicate the source of the biomarker genes. The small size nodes are the connection genes linking these biomarker genes on the BioGrid network.

In addition, we conducted the survival analysis and metastasis comparison using the selected network biomarker genes (see **Fig. 2)**. As seen, the selected biomarkers can divide the patients **into** two sub-groups with distinct survival rates in both G3 (p-value = 0.012) MBs. However, it is failed to identify these subgroups using gene expression signature of all genes in both G3 (p-value = 0.14). In addition, for the metastasis, a sub-group in G3 MBs is identified with less than 1/3 (9 vs. 31) metastasis rate vs. 50% metastasis rate (34/35), which is more separated than the subgroups (16/21 vs 27/45) obtained using the all gene signature. As seen, the network signature can also identify the sub-groups with more females in subgroup 1 vs more males in subgroup 2. The results indicate the network biomarker genes have the capacity to separate these phenotypes. To further validate the network biomarker genes, we also randomly selected sets of genes, with the same size of the network signature, for the survival, metastasis and sex subtyping analyses, and repeated these analyses 10,000 times. The p-values observed using on the network signatures, compared with using all the genes, were 0.027, 0.015 and 0.0002 respectively, which indicate the significance of the network biomarker genes.

**Figure 2.**
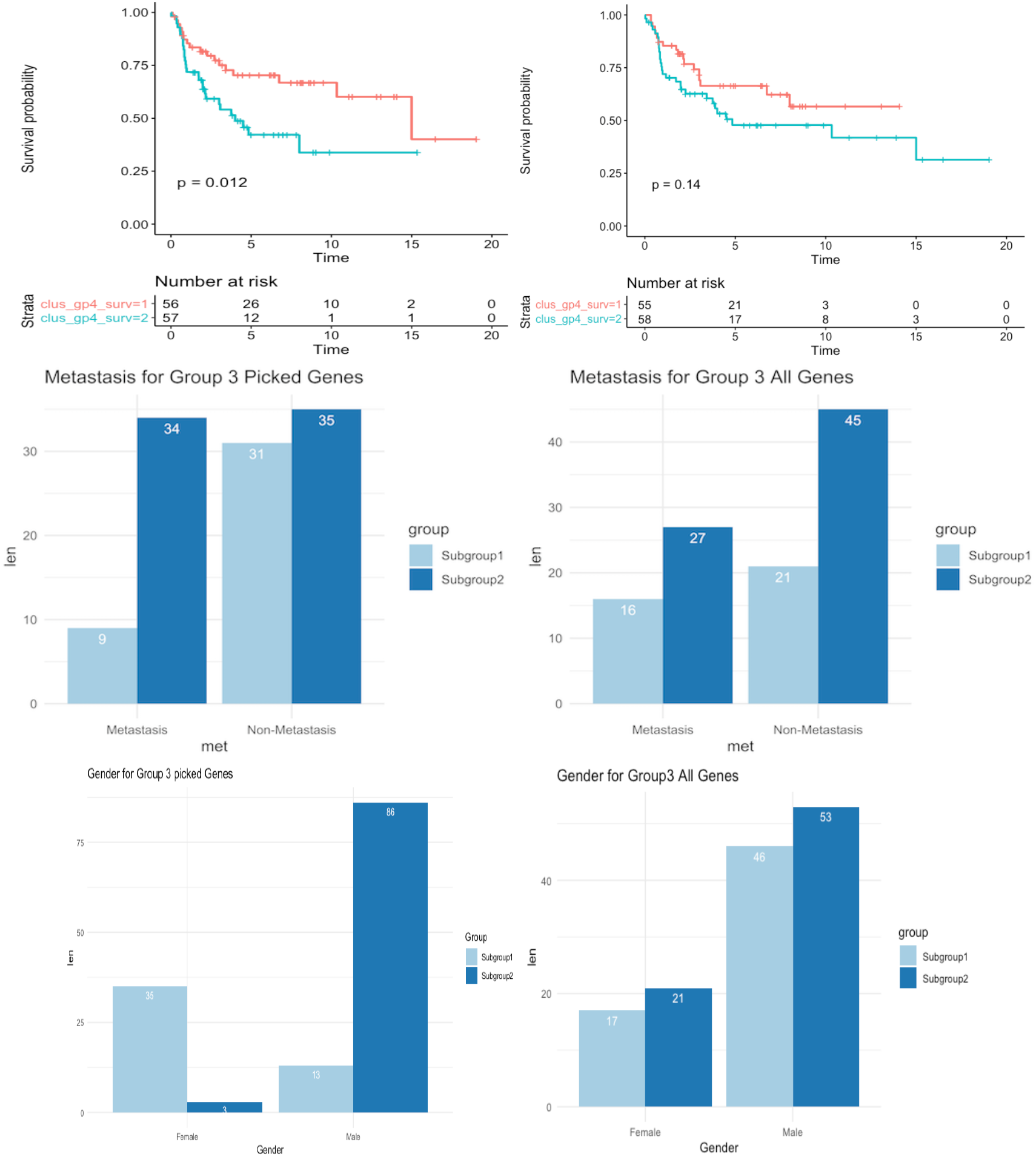
Survival analysis (**upper-panel**), Metastasis difference (**middle-panel**), and gender difference **(bottom-panel)** of G3 MBs using selected network biomarkers (**left-panel**) and all genes (**right-panel**).

### 3.3 Identify drugs potentially targeting on the network biomarkers

We made use of the integrative signaling network as a network signature to predict drugs that can potentially inhibit/perturb the targets on the signaling network. First, we identified drugs that can directly inhibit genes on the signaling network by using the drug target information derived from DrugBank database (version 5.1.4, released 2019-07-02). **Fig. 3** shows the FDA-approved drugs that directly inhibit the genes on the signaling networks of G3 (12 total drugs). In addition, we identified drugs that can potentially inhibit the network genes indirectly by using the CMAP database, which provides the drug search tool in their website (clue.io). We selected all the drugs and investigational compounds that have a score <= −80, which means the drugs can potentially reverse the gene expression of the genes on the signaling network. To further investigate the mechanism of these drugs based on their intended targets, we clustered the selected drugs based on their targets. **Fig. 4** shows the clusters of FDA approved drugs and all CMAP drugs and compounds for G3 MBs. As can be seen, some drugs target a DNA topoisomerase, MAPK, PI3K, and HDAC, which have been identified by high throughput screening to be effective in inhibiting MYC-driven Medulloblastoma cells^6^. Especially, the PI3K and HDAC inhibitors synergistically inhibit the growth of MYC-driven MB cells. Also, drugs that target genes such as GABA and FGF have also been reported to be effective for MB treatment ^53,54^.

**Figure 3.**
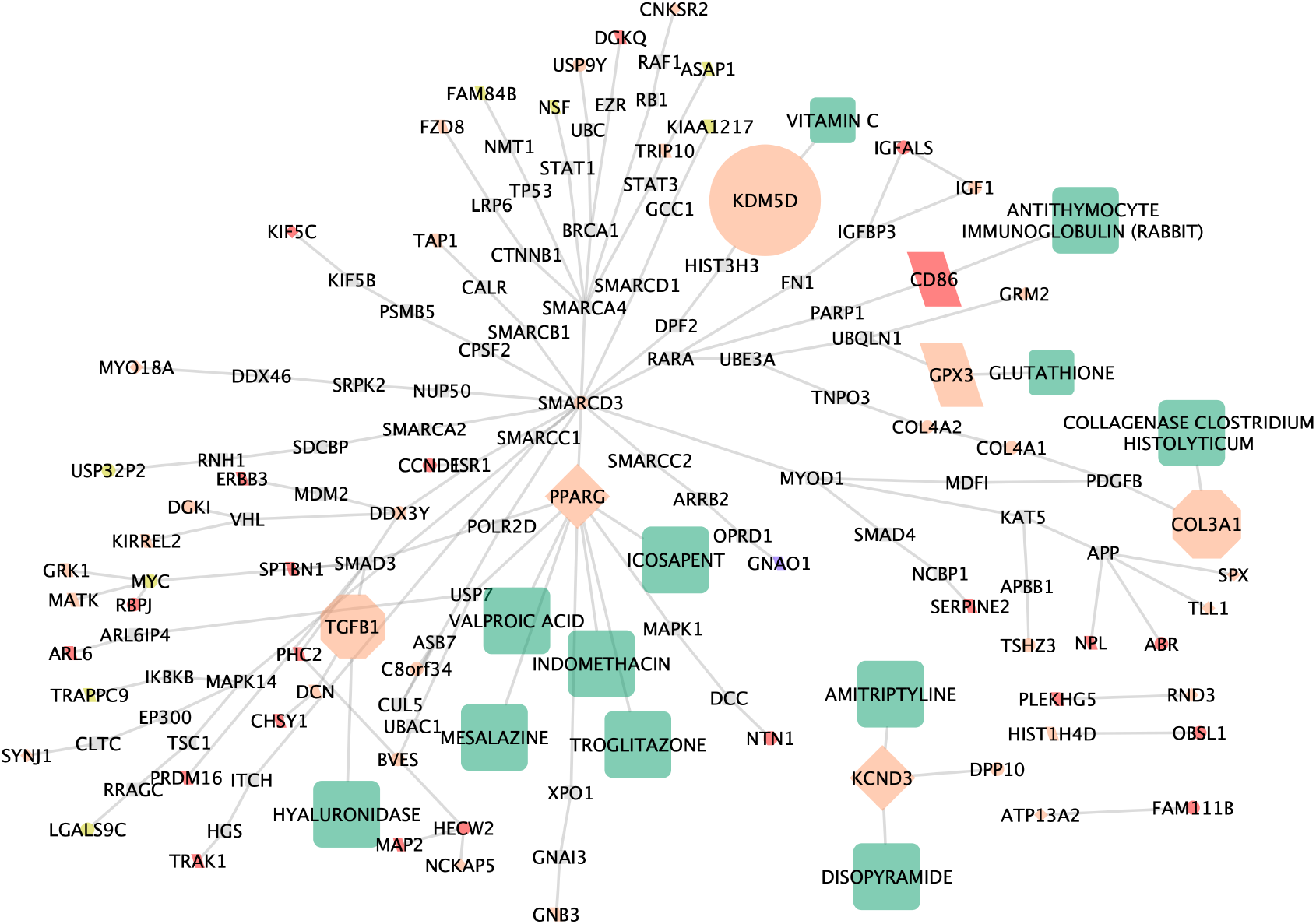
FDA approved drugs targeting on the network biomarker genes of G3 MBs.

**Figure 4.**
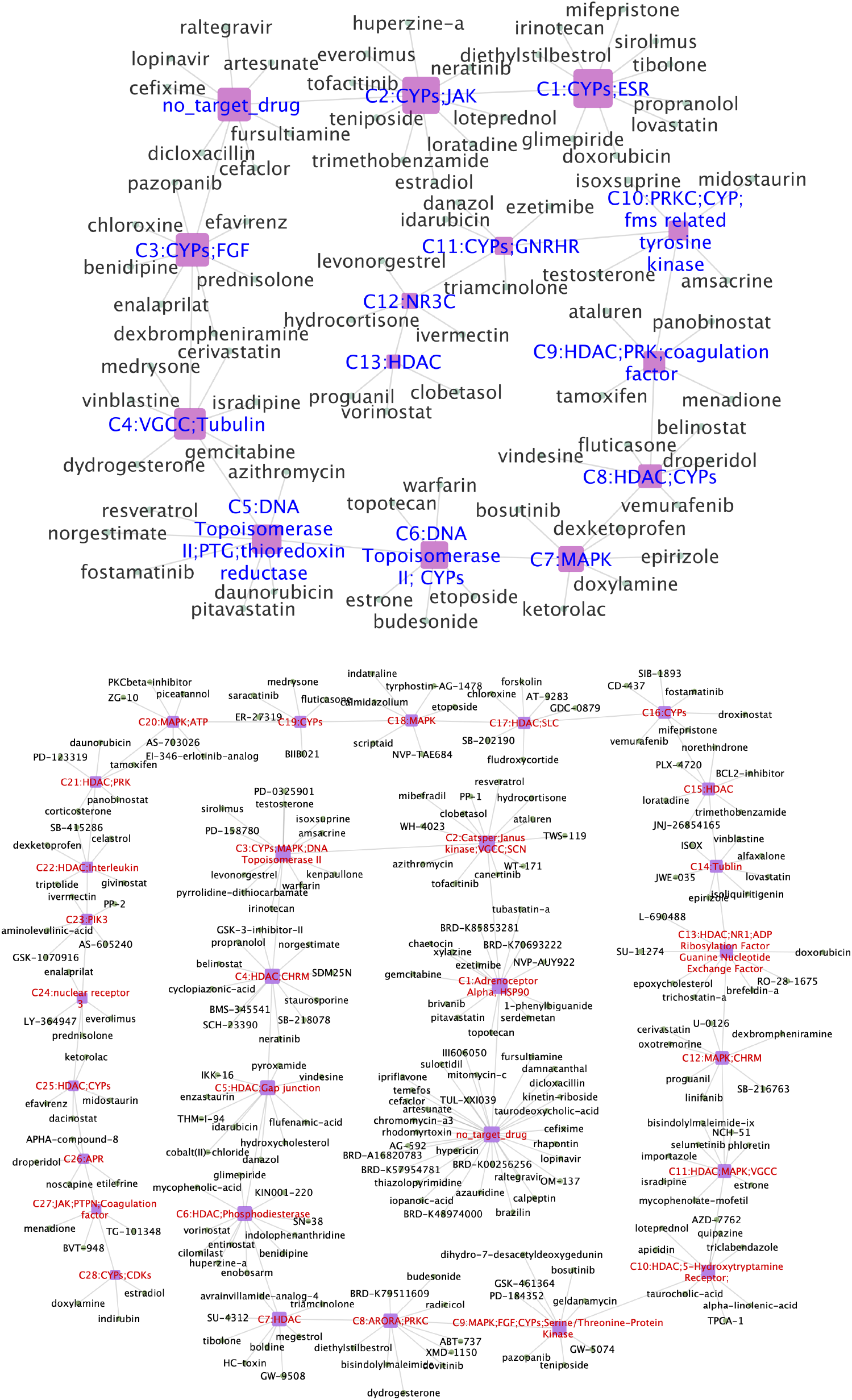
Drug clusters of FDA approved drugs (upper) and all CMAP drugs and compounds that can potentially inhibit the network biomarker genes of G3 MBs.

## 4. Summary and discussion

Medulloblastoma (MB) is the most common malignant brain tumor in infants and children. Among the four subtypes of MB, G3 and G4 MBs have the most ambiguous signaling mechanisms and poor outcomes as compared to WNT and SHH subtypes. Also, there is no effective targeted therapy for G3. Using the available multi-omics data (gene expression, methylation and copy number/SNP) from MB patients, we identified potential biomarkers that were associated with three phenotypes: poor survival rate, high metastasis rate, and sex difference (much higher prevalence rate in boys than girls). Through the integrative analysis, we identified a set of biomarker genes that are associated with worse survival rate, metastasis and gender difference respectively. Moreover, the signaling network signature was obtained by linking these selected biomarkers in the BioGRID protein-protein interactome. We then identified drugs that can potentially inhibit the signaling network using drug target information and reverse gene signature in the CMAP database. Many of these targets and drugs have been reported to be related to MBs, thereby validating our approach.

While challenging, it is important to uncover the causal signaling cascades that regulate the three phenotypes and to discover novel and effective targeted therapies for G3 MB treatment. This study is exploratory research aimed to identify the potential biomarker genes associated with the three phenotypes. The causal relationships among these biomarkers remain unknown. In the future work, we will evaluate these biomarkers, drugs and drug combinations using in vitro G3 MB cell line models or patient-derived xenograft (PDX) models.

## Notes

### Competing Interest Statement

The authors have declared no competing interest.

